# Stony coral tissue loss disease outbreak altered reef communities of Carrie Bow Cay, Belize

**DOI:** 10.64898/2026.09.09.750164

**Authors:** Leah M Harper, Brooke Sienkiewicz, Logan Laurent, Tristin Henson, Christian Flaherty, Viviana D Bravo, Scott Jones, Greta Aeby, Valerie Paul, Sarah Gignoux-Wolfsohn

## Abstract

Stony coral tissue loss disease (SCTLD) has caused one of the most damaging coral epizootics on record, impacting about half of coral species across most of the reef systems of the Caribbean and Florida. To evaluate its impact on the coral reefs surrounding Carrie Bow Cay, Belize, we monitored seven fixed transects before and at two time points during the outbreak using three different methods. We estimated species-specific mortality rates by fate tracking tagged colonies, quantified reef-wide lesion prevalence and resulting changes to colony density and community composition using in situ transect surveys, and finally calculated changes to total coral cover using benthic photoquadrats. SCTLD quickly led to mortality of infected corals. Between October 2019 and May 2022, 32% of the 102 fate-tracked coral colonies had developed lesions consistent with SCTLD and an additional 30% had suffered complete mortality likely due to SCTLD. By December of 2022, 21% of the 33 colonies that had been observed with lesions had died. We documented patterns of species susceptibility largely confirming previous reports in other locations. We documented sharp abundance declines in several species, including the total loss of *Meandrina meandrites* from survey transects, leading to a significant shift in community composition. Cover of several species declined between October 2019 and May 2022, but only *Siderastrea siderea* experienced sustained declines into December of 2022. Overall cover of scleractinian corals decreased from 13.2 to 7.9% during the monitoring period. The ecological impacts of SCTLD in the area surrounding Carrie Bow Cay are consistent with reports from elsewhere in the Caribbean, underscoring the magnitude of colony density and coral cover loss caused by SCTLD across the region.

## 1. Introduction

The coral reef ecosystems of the Caribbean have been burdened by diseases affecting reef-building corals for at least half a century (Garrett and Ducklow 1975; Gladfelter 1982; Edmunds 1991; Richardson 1998), with reports of disease outbreaks becoming more frequent and widespread over time (Lafferty et al. 2004; Sokolow 2009; Tracy et al. 2019). Most recently, stony coral tissue loss disease (SCTLD, (Hawthorn et al. 2024)) has emerged as an epizootic widely considered to be the most damaging coral disease yet identified (Papke et al. 2024). SCTLD affects approximately half of all Caribbean coral species with variable rapidity and severity. For the most susceptible species the disease can kill entire coral colonies within weeks after lesion appearance (Aeby et al. 2019; Estrada-Saldívar et al. 2020; Meiling et al. 2021), contributing to local extirpations of highly susceptible species in several locations (Alvarez-Filip et al. 2019; Neely et al. 2021a; Jones et al. 2021; Brandt et al. 2021; Buenrostro-Muñoz et al. 2026; Álvarez-Filip et al. 2026).

After a decade of research on SCTLD, many uncertainties about the etiology, transmission, and ecological impacts of the disease persist. Notably, as with most coral diseases, the causative agent of the disease remains unknown (Richardson 1998; Rogers 2010; Ushijima et al. 2020). In addition, discrepancies across lab and field conditions as well as inconsistencies in the manifestation of disease signs have resulted in some disagreement on how susceptibility to SCTLD varies across species. Specifically, the susceptibility of the common genus *Porites* differs across lab exposure experiments (Aeby et al. 2019; Meiling et al. 2021; Papke et al. 2024) and the genus *Agaricia*, while generally found with very low or no disease prevalence (Alvarez-Filip et al. 2019; Hayes et al. 2022a), has been reported affected in the USVI (Brandt et al. 2021) and has not been tested in the lab (Papke et al. 2024). In the field, *Siderastrea siderea* often exhibits atypical dark or purple lesions that progress slowly sometimes in addition to the white acute and subacute SCTLD lesions characteristic of SCTLD (Hawthorn et al. 2024). These chronic lesions have been suggested to potentially be caused by a different disease entirely (Aeby et al. 2025). Determining in situ variation in the effects of SCTLD across taxa, space, and time is crucial to understanding and predicting its current and future effects on Caribbean coral reef ecosystems. This information can only be obtained through clear, quantitative reports of disease severity and its impact on coral abundance or cover (and mortality). Common methods for quantifying disease severity include fate-tracking of individual colonies (Precht et al. 2016; Rippe et al. 2019; Sharp et al. 2020; Kolodziej et al. 2021) and transect surveys enumerating colony density and lesion prevalence (Alvarez-Filip et al. 2019; Estrada-Saldívar et al. 2020; Croquer et al. 2022; Hayes et al. 2022a). Ecosystem-level effects of disease outbreaks are often measured through changes in cover of scleractinian corals, an essential ocean variable (Miloslavich et al. 2018) with broad utility for tracking and interpreting the condition of coral reef habitats. Any observed ‘phase shifts,’ defined as rapid, persistent changes in benthic community structures can potentially signal long-term changes in ecosystem function (Crisp et al. 2022).

Since the first account of SCTLD in Florida in 2014 (Precht et al. 2016; Walton et al. 2018), it has spread throughout the Caribbean (Estrada-Saldívar et al. 2020; Heres et al. 2021; Williams et al. 2021; Dahlgren et al. 2021; Brandt et al. 2021; Alvarez-Filip et al. 2022; Lee Hing et al. 2022; Truc et al. 2023), leading to widespread shifts in benthic community composition, loss of live coral cover (Heres et al. 2021; Brandt et al. 2021; Jones et al. 2022), and resulting declines in ecosystem functions including calcium carbonate accretion and habitat complexity (Alvarez-Filip et al. 2022; Swaminathan et al. 2024). However, since reef communities and environmental conditions vary across Caribbean reefs, documenting the impacts of SCTLD in disparate locations can help us to understand both region-wide effects and factors that influence disease progression. Along the Mesoamerican Reef (MAR), the world’s second-largest barrier reef, SCTLD was first reported in summer 2018 in Quintana Roo, Mexico (Kramer et al. 2019). The leading edge of the disease outbreak reached northern Belize in June 2019. While spread over the following two years was slow, potentially due to prevailing northern currents, SCTLD impacted nearly all of the MAR by the summer of 2021 (Lee Hing et al. 2022).

Here, we report the effects of SCTLD on reefs surrounding Carrie Bow Cay in Belize using a suite of monitoring metrics. To understand the ecosystem changes that are attributable to the disease outbreak, rather than other stressors such as recurrent thermal anomalies, we established a baseline fate-tracking and monitoring program before SCTLD reached the area. Both before and after the arrival of SCTLD, we conducted in situ and photographic coral monitoring to measure coral density, disease prevalence, and cover through time. These data add to the growing body of literature quantifying the impacts of SCTLD on Caribbean reef systems.

## 2. Methods

### 2.1 Study sites

Seven study sites were selected from a suite of permanent transects established by the Smithsonian’s MarineGEO and Caribbean Coral Reef Ecosystems programs close to Carrie Bow Cay, Belize (Fig. 1). Four sites on the forereef were chosen, ranging in depth from 8.5-11m, along with three patch reef sites in the lagoon with depths of approximately 4-4.5m (Table S1). Sites were chosen for accessibility, sufficient spatial area to establish transects, and sufficient density of target species close to the transect. Transect start and end points are fixed with permanent markers.

**Figure 1.**
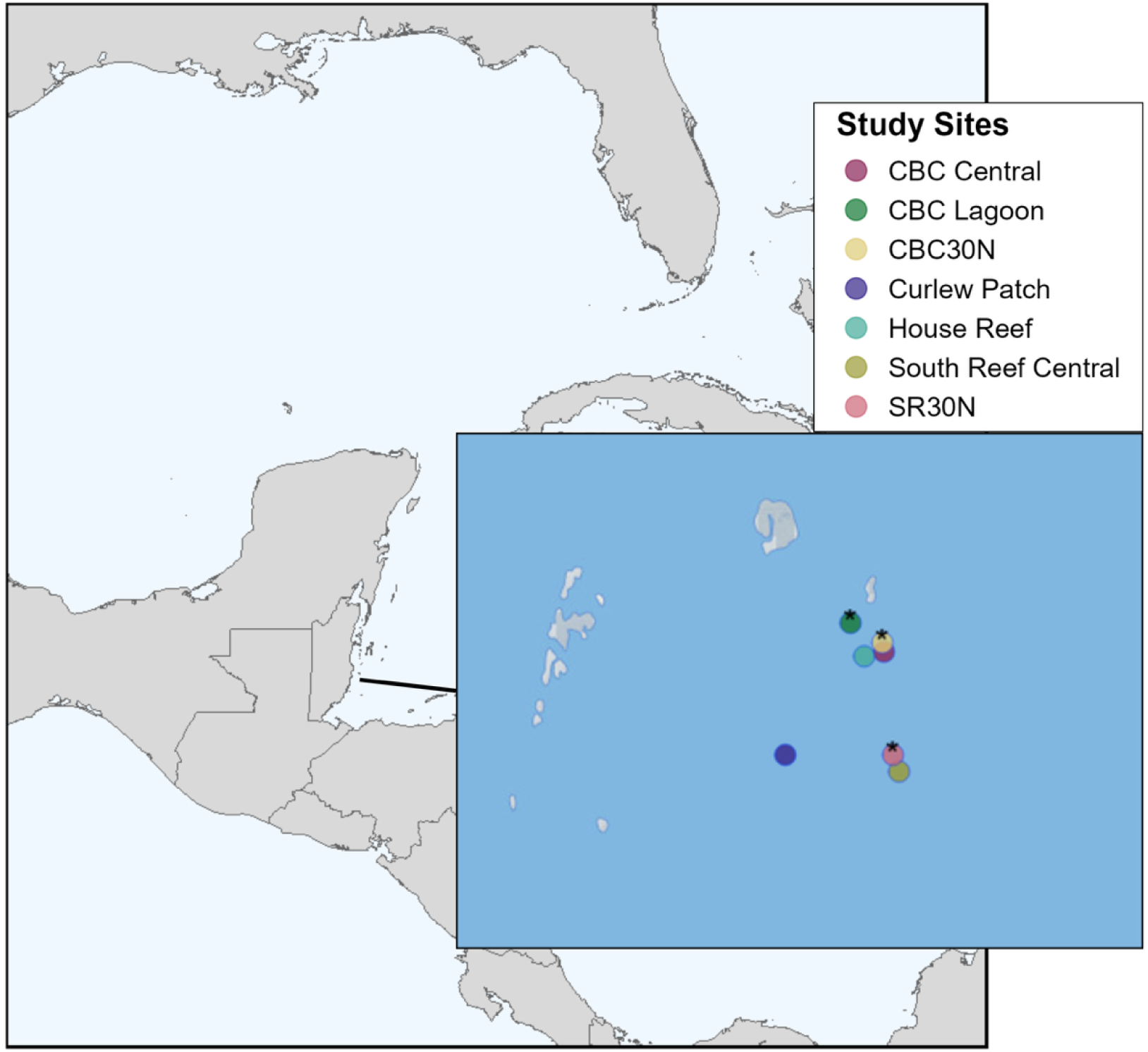
Study sites surrounding Carrie Bow Cay, Belize. Map produced using Leaflet.

### 2.2 Colony fate-tracking

In June 2019 we selected 81 colonies for fate tracking across three of the permanent transects. Along each transect, the healthiest-appearing colonies in reasonable proximity to the transect line (<10m), with no signs of active tissue loss, color loss, or discoloration, were chosen for each of the following species: *Montastrastraea cavernosa* (n=24), *Siderastrea siderea* (n=27), *Porites astreoides* (n=17), and *Meandrina meandrites* (n=13). In October 2019, stony coral tissue loss disease had still not reached Carrie Bow Cay, and we added 21 healthy-appearing colonies of *Pseudodiploria strigosa*. Colonies were photographed, tagged, and mapped.

Stony coral tissue loss disease reportedly reached Carrie Bow Cay in July 2021, while the station was closed due to the Covid-19 pandemic (Kramer et al. 2019; Carne 2021). We visited upon its reopening in May 2022, located the tagged colonies, photographed them (Fig. 2), and documented signs of disease. For all species other than *S. siderea*, colonies were classified as “diseased” (active tissue loss) when they presented with bare white skeleton next to live tissue along one or more focal lesions (Fig. 2A, B and C). For *S. siderea*, diseased colonies displayed tissue loss in a diffuse distribution (Raymundo et al. 2008) Fig, 2D). Colonies were considered “dead” when they had complete mortality, meaning no living tissue (diseased or healthy) remained on the skeleton (Fig. 2A). This fate-tracking survey was repeated in December 2022.

**Figure 2.**
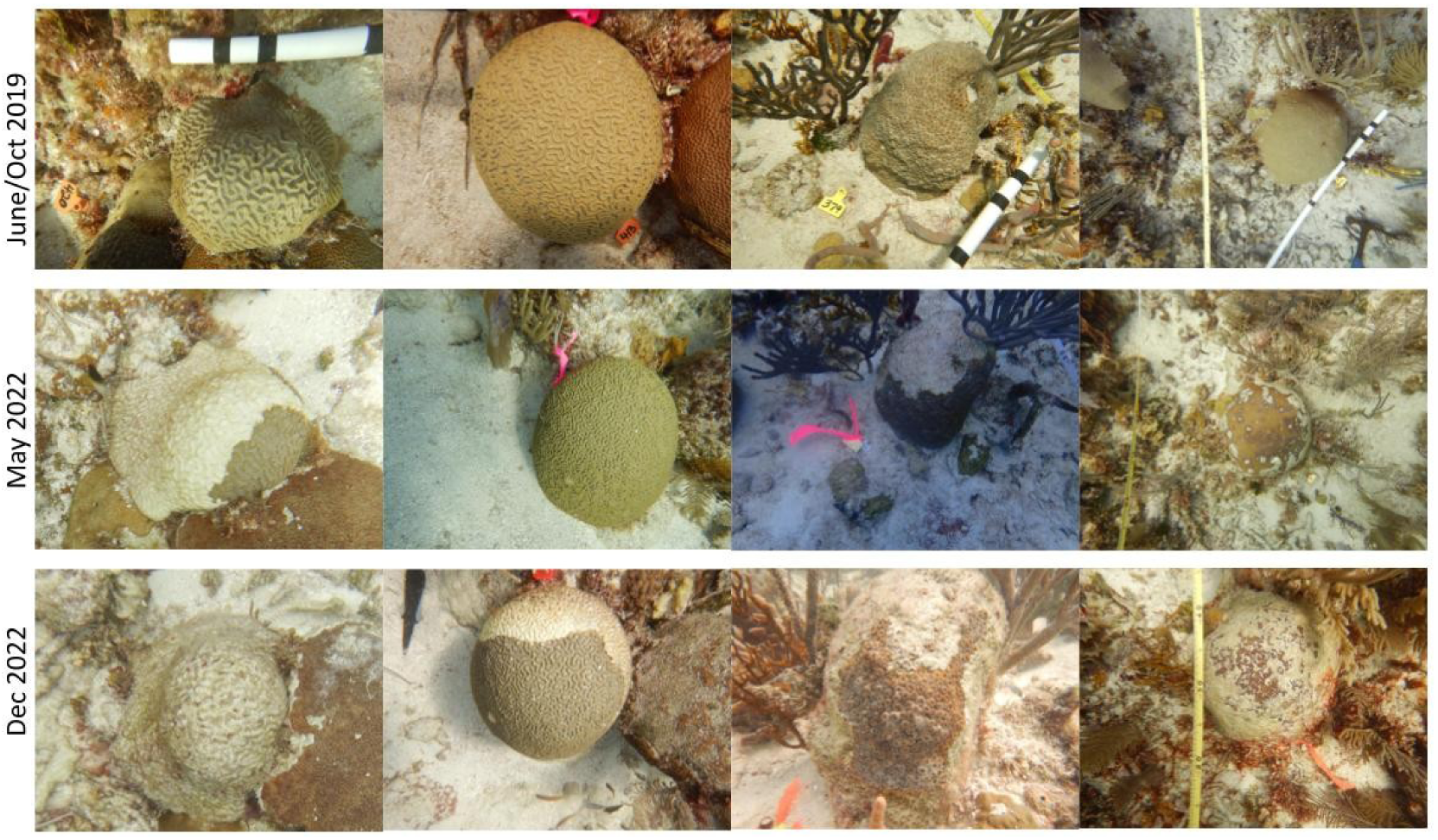
Four example coral colonies tracked over time. All four were healthy in 2019 and subsequently developed disease signs and either partial or complete mortality.

Several tagged colonies were completely dead by the second timepoint (May 2022), including all of the *M. meandrites*. While it is impossible to know the exact cause of death, we make the reasonable assumption (see: (Precht et al. 2016; Rippe et al. 2019; Estrada-Saldívar et al. 2020) for previous examples) that these colonies died from SCTLD. We do not include these colonies in calculations of tissue loss prevalence at a given timepoint but do include them in calculations of “cumulative prevalence,” *i.e.,* prevalence across the entire study period. We therefore calculate tissue loss prevalence as:

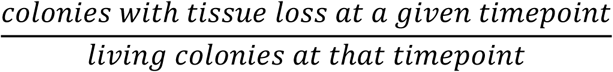

and cumulative prevalence as:

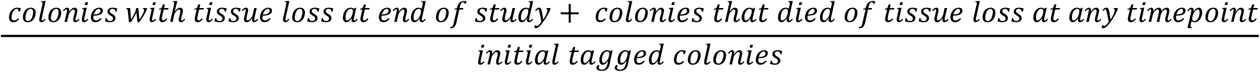

We estimated changes in partial mortality of individual colonies over time using the photographs of colonies at each time point, with the first photo taken at initial tagging as a reference. Percent live tissue was estimated in 5% increments by two independent observers and averaged. Percent change was then calculated as (final % live tissue - initial % live tissue / initial). To quantify relative rates of tissue loss caused by the SCTLD outbreak for each tagged colony, the rate of change in percent live tissue for May - December 2022 was calculated as (percent change / # weeks between timepoints). Colonies that had already died by May 2022 were therefore excluded, including all *M. meandrites.* Initial maximum colony diameter was not correlated with rate of tissue loss (Spearman’s ρ = 0.126, p-value = 0.217). We recognize however that our small sample size may have limited a clear relationship between size and tissue loss, and since we did not measure area of live tissue or lesions, these serve as relative rates to compare tissue loss between species.

### 2.3 In situ coral surveys

We used the MarineGEO protocol to survey coral demographics and conditions at three timepoints (Table S1), (MarineGEO 2026). Five out of the seven sites were surveyed in October 2019, with the remaining two surveyed in January 2020 and all seven were surveyed again in May and December 2022. In brief, all living corals within a single 30m long x 1m wide belt transect at each site were counted, identified to species in most instances, and placed into size classes. Due to the difficulty of consistent in situ identification, *Orbicella faveolata*, *O. franksi*, and *O*. *annularis* were grouped into an Orbicella species complex (referred to as “*Orbicella* spp.”); *Porites porites*, *P. divericata*, and *P. furcata* were grouped into a *Porites porites* species complex (referred to as “*Porites porites*”); and *Agaricia agaricites* and *A. humilis* were grouped into an *Agaricia agaricites* species complex (referred to as “*Agaricia agaricites*”). Health status of colonies was assessed as described above for tagged colonies. Disease prevalence was calculated at a given timepoint for each species and transect by dividing the number of colonies with lesions by the total number of colonies of each species at that timepoint.

### 2.4 Benthic photoquadrats

At the time of each demographic survey, 16-21 benthic photoquadrats were captured for each transect. Photos were taken approximately 1m above the benthos using a standard distance framer (Fig. S1) approximately every 2m along the 30m transect line. Within each photo, 40 random points were assigned using CoralNet’s online platform for a total of 640-840 points per transect per survey. Each point was annotated to the species level for scleractinian corals. Non-coral points were annotated to benthic functional group (e.g. octocoral, macroalgae, turf algae, sponge).

### 2.5 Statistical methods

To test for change in species-level community composition of scleractinians recorded by in situ transect surveys through time and across locations, we used the ‘adonis’ function in the ‘vegan’ package to perform a PERMANOVA with time and transect as factors, and visualized Bray-Curtis dissimilarity between sites and survey timepoints using non-metric multidimensional scaling (NMDS). We performed this PERMANOVA test both with and without juvenile corals (<4 cm). We investigated changes in transect-level abundances of coral species across time and location by using the ‘GLMMadaptive’ package in R to fit a zero-inflated negative binomial mixed effects model, with the count of all colonies, including juveniles, within the 30m^2^ transect area, as the response variable. We included survey timepoint, species, and their interaction as fixed effects and transect as a random effect, specifying species as the zero-inflated fixed effect and transect as a zero-inflated random effect because some species were absent from some transects even before SCTLD onset. We excluded all species with fewer than eight total observations to further address zero inflation (Table S2). The model failed to converge, so we further simplified the parameters by dropping *Madracis decactis* and *Helioseris cucullata*, which had 26 and 16 total observations, respectively, but were not species of interest in our investigation of SCTLD due to their combination of scarcity and low susceptibility. Results with and without these species included in the model were similar (both main effects and their interaction were significant); however, their removal allowed the model to converge. We report the results of the converged model below. We also tested the change in tissue loss prevalence over time. We limited this analysis to those species that experienced at least one incidence of tissue loss during the study period. We used the ‘GLMMadaptive’ package to fit a beta binomial mixed model with the counts of healthy corals and corals with tissue loss as the response variable, survey timepoint and species as fixed effects, and transect as a random effect. We attempted to fit this disease prevalence model with the survey × species interaction, but the model failed to converge despite our optimization strategies (adjusting starting values, increasing iterations). The failure to converge indicates the data are insufficient to reliably estimate interaction effects, so we present results from a main effects model.

For both the colony abundance and tissue loss prevalence models, we used Wald tests to determine significance of survey timepoint, species, and for colony abundance their interaction, and used the package ‘emmeans’ to obtain pairwise significant differences across time points and separately, among species, at a threshold p < 0.05.

To test whether cover of benthic functional groups (scleractinians, octocorals, macroalgae, and turf algae) calculated from photoquadrats differed across the three study time points, we pooled all classified points across the 16-21 photos from each survey as unit of replication (e.g. the combination of site and time point) and calculated proportion cover as the number of points of each benthic component divided by total number of points, and percent cover as that proportion multiplied by 100. We used a binomial generalized linear mixed model in the ‘lme4’ R package, in which time point, functional group identity, and their interaction were fixed effects, site was a random effect, and proportion cover (weighted by the number of total available classified points) was the response variable. We used the package ‘emmeans’ to obtain pairwise significant differences at a threshold p < 0.05. All points identified as scleractinian coral were included in the benthic functional group analysis, regardless of species. This process was repeated to identify species-specific percent cover differences over time using the same list of species analyzed for density change with percent cover as a response and time point, species, and their interaction included as fixed effects, but we specified a BOBYQA (Bound Optimization BY Quadratic Approximation) optimizer within the generalized linear mixed models to improve convergence.

A non-parametric Kruskall-Wallis test was used to compare rates of tissue loss from May to December 2022 between tagged *M. cavernosa*, *S. siderea*, *P. astreoides*, and *P. strigosa* because the data did not meet parametric assumptions. Only coral colonies observed with disease signs in May or December 2022 were included; colonies that were completely dead in May 2022 and colonies that remained healthy throughout the study were excluded. Dunn’s test with a Bonferroni correction was used to evaluate post-hoc pairwise comparisons. These were completed with the ‘rstatix’ package in R.

## 3. Results

### 3.1 SCTLD is fatal with progression speed varying by species

While all tagged coral colonies were healthy upon first visit in June or October 2019, 31 of the initial 102 were already dead by May 2022, after SCTLD had been reported in the vicinity approximately 10 months prior (Fig. 3). Once corals displayed disease signs, they usually did not recover. Thirty-three coral colonies displayed active disease signs in May 2022. Of these, 7 were dead by December 2022. Only 5 colonies seemed to recover, meaning there were no active disease signs in December 2022, but see below for species-specific considerations.

**Figure 3.**
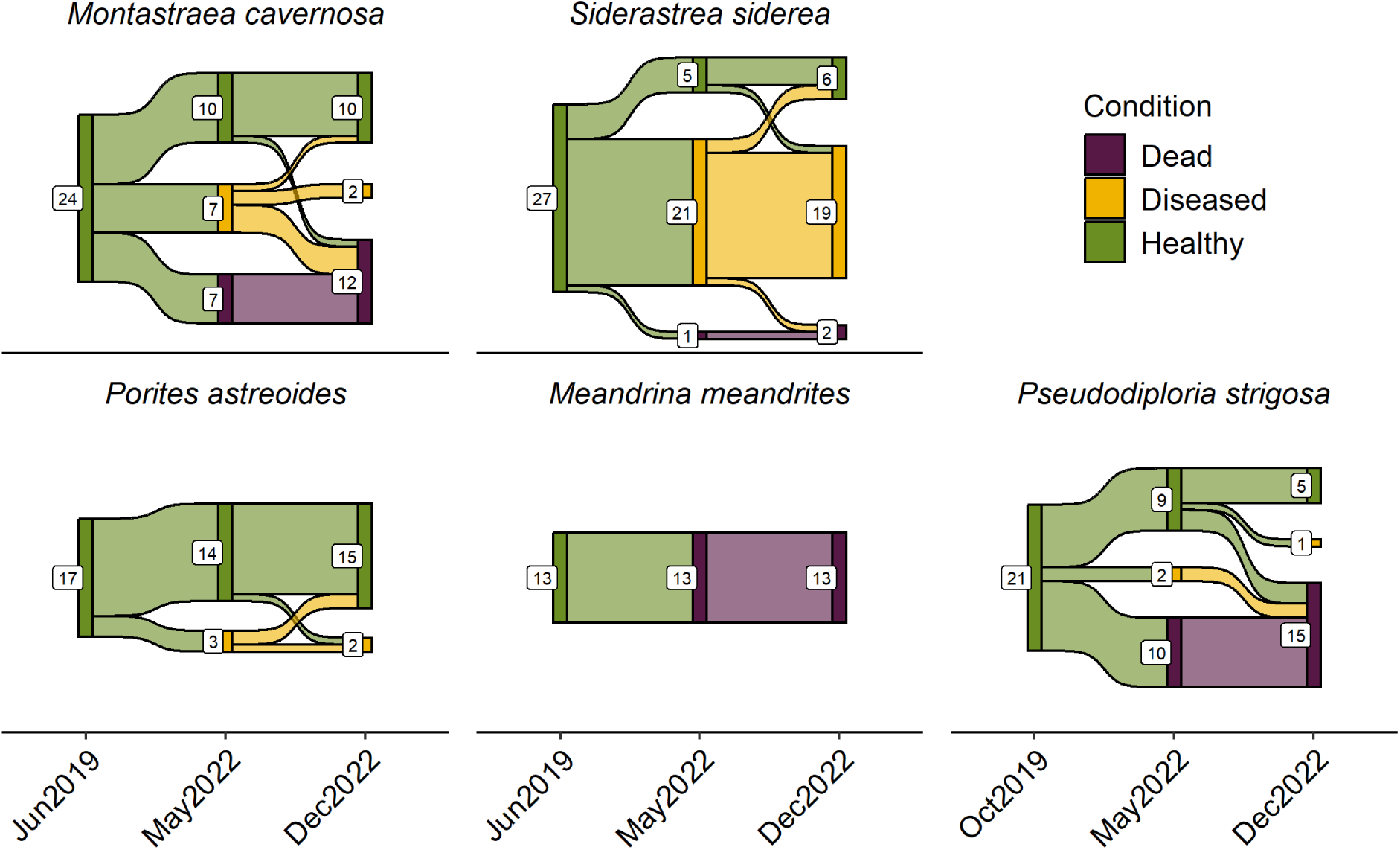
Sankey plot showing fate of individual tagged corals at three timepoints. Numbers indicate the number of colonies with a given condition at that time, individual colonies are connected by lines.

The highest prevalence and (inferred) fastest disease progression among tagged colonies occurred in *M. meandrites* – all 13 colonies tagged in 2019 were completely dead by May 2022 (100% mortality). *P. strigosa* had the next highest prevalence and fastest progression: of the 21 colonies tagged in 2019, 10 were completely dead (48% mortality) and 2 diseased by May 2022. By December 2022, only 5 tagged *P. strigosa* were still healthy, resulting in 76% cumulative prevalence and 71% mortality. *P. strigosa* had the highest relative rate of tissue loss May - December 2022 (Fig. 4; mean = -3.2% week^-1^ SD 0.91) which was significantly higher than *P. astreoides* (p = 0.048) and *S. siderea* (p = 0.003). The disease progressed more slowly in *M. cavernosa*, with 7 of the initial 24 tagged corals dead and an additional 7 diseased by May 2022 (42% cumulative prevalence). Eight of the remaining healthy *M. cavernosa* were still healthy in December 2022, while one had completely died and one previously diseased *M. cavernosa* appeared to recover by December (Fig. S2). Disease progression was slowest in *S. siderea*, although by the end of the study period prevalence was high (85% cumulative prevalence). The relative rate of tissue loss in this species was low (mean = -0.74% week^-1^ SD 0.89) between May and December 2022, and was significantly lower than *P. strigosa* and *M. cavernosa* (p = 0.005). Only one *S. siderea* was dead in May 2022, while 21 colonies were diseased (81% prevalence). Of these, only one had died completely by December 2022. Two colonies that appeared diseased in May were apparently healthy by December. Disease signs for *S. siderea* presented differently than all other species, with multifocal lesions covered by spider-web like mucus (figures 2 and S2). The least affected species was *P. astreoides,* with only 3 out of 17 tagged colonies displaying tissue loss in May 2022, 2 of which had recovered by December 2022. We did not document any *P. astreoides* complete mortalities among tracked colonies.

**Figure 4.**
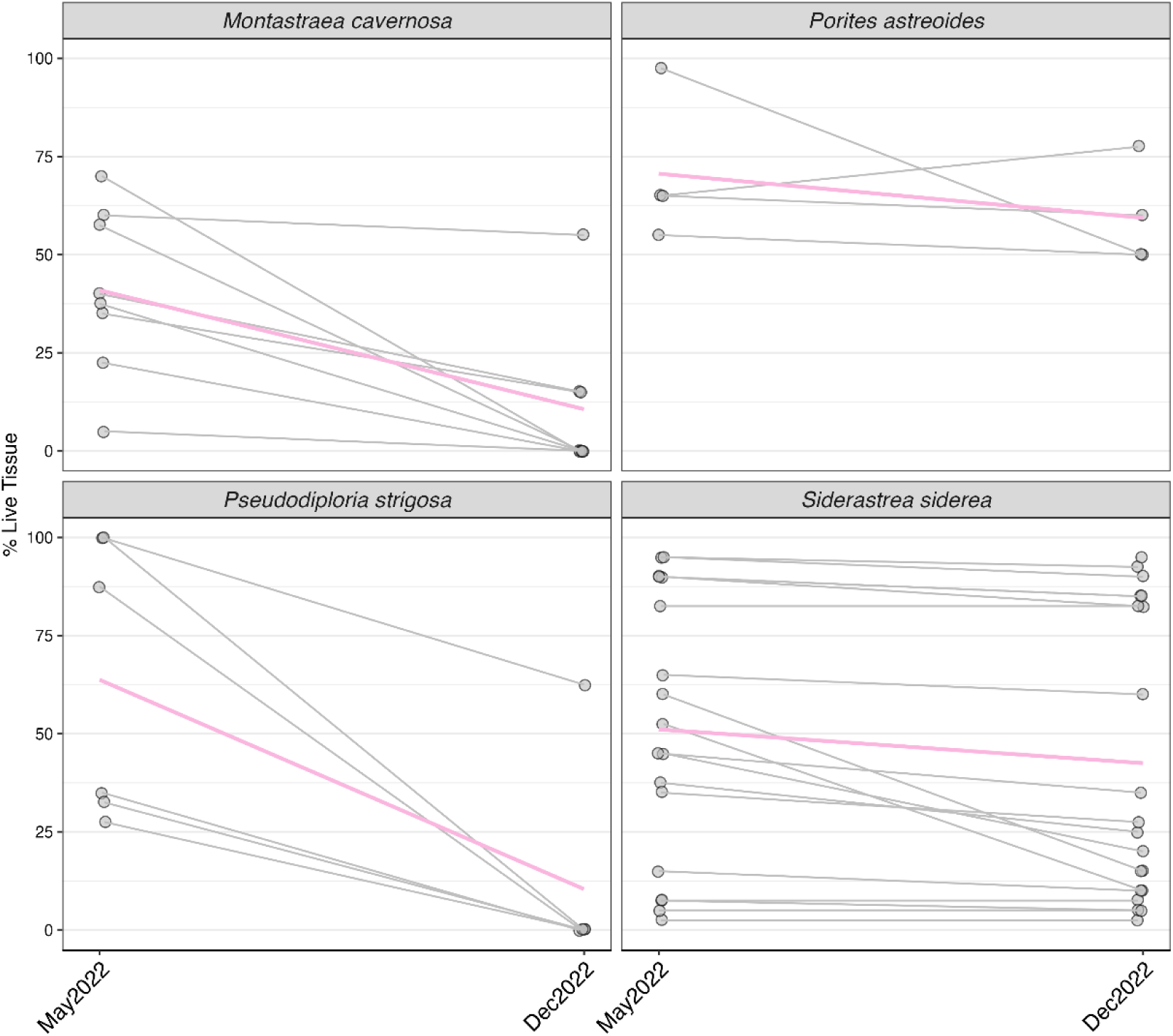
Percent live tissue (relative to initial tagging) of tagged colonies that displayed signs of tissue loss between May and- December 2022. The pink line is the mean percent live tissue for each species, and grey lines show individual colonies over time.

### 3.2 Prevalence and incidence of SCTLD varies across species

At the time of the initial in situ surveys (October 2019/January 2020), tissue loss lesions were rare (0.3% of all surveyed corals) at all of our transects and were only reported in agaricids and poritids (Fig. 5, Fig S4, Table S2). Overall lesion prevalence across all species increased significantly to 6.8% in May 2022 (Fig. 5; GLMM, Chisq = 58.087, DF = 2, p < 0.001, Table S3). Prevalence of tissue loss differed significantly by species (GLMM, Chisq = 137.383, df = 11, p <0.001), with agaricids and poritids generally exhibiting fewer lesions than those species that were lesion-free at the start of the study (Fig. 5, Table S3). *P. strigosa* had the highest prevalence of active SCTLD-lesions across all sites in May 2022, with 30.4% lesion prevalence across 23 corals at 5 sites. In December 2022, *S. siderea* had the highest prevalence with 31.9% prevalence across 129 corals at all 7 sites. We also found that 4 of the 6 *E. fastigiata* found in May 2022 were infected (66.7%). While we recorded high prevalence at sites with few *M. cavernosa* (e.g., the one *M. cavernosa* at CBC Central was diseased in May 2022), across all sites *M. cavernosa* had lower prevalence of tissue loss (24.1%). The *Orbicella* species complex had the lowest lesion prevalences of the affected coral species (17.4% peak prevalence across sites in May 2022). While *S. siderea* had low overall prevalence in May 2022 (18.9%), it was the only species where prevalence increased between May and December 2022 (to 31.9%) across all sites. *P. strigosa* was the only species to maintain its prevalence through December 2022 (31.3%) while all other species saw reduced tissue loss prevalence over time.

**Figure 5.**
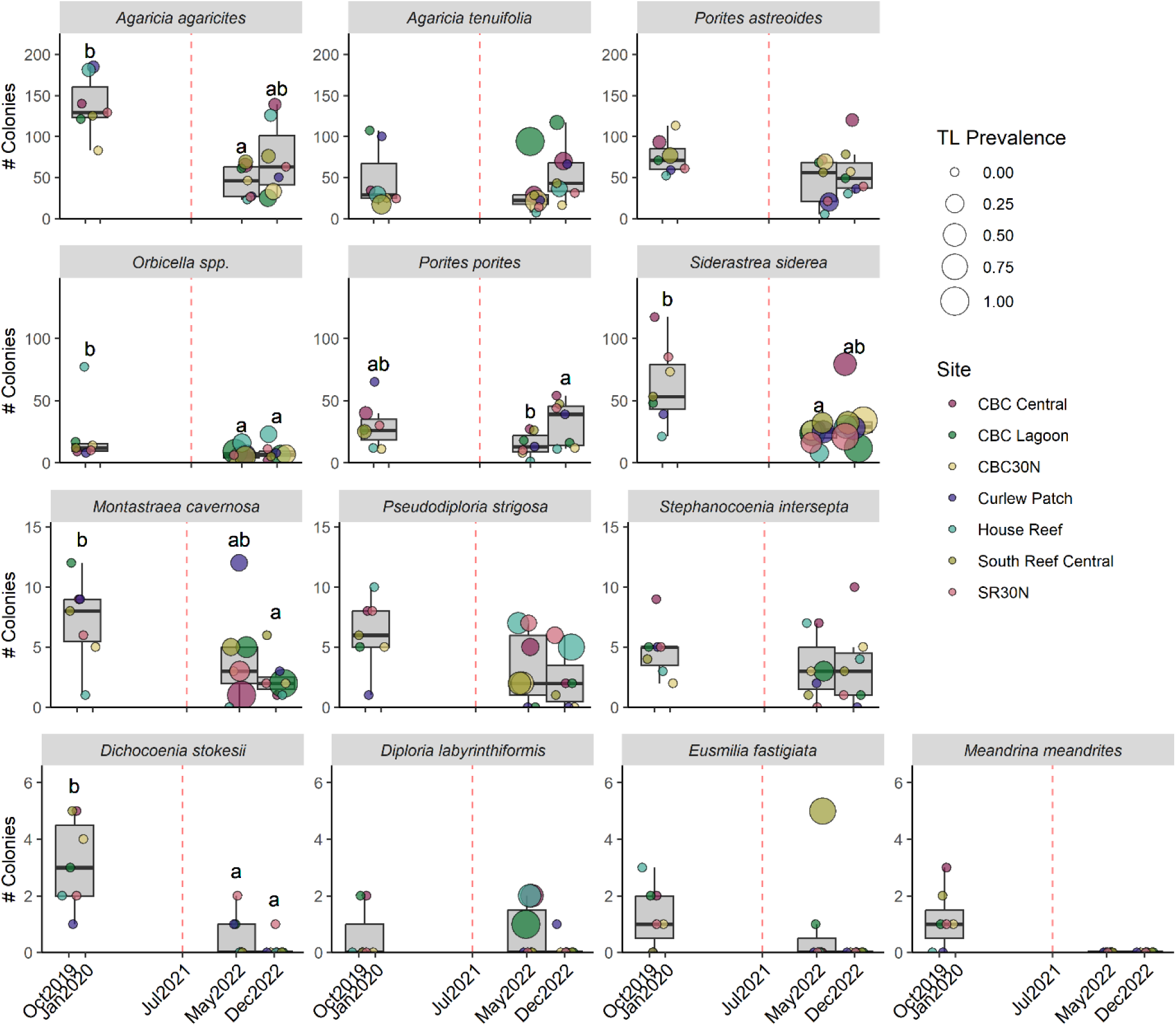
Colony abundance recorded during in situ surveys for select species along a 30m x 1m transect at each site over time. All species presented here were included in the glmm, data for all species is presented in Table S1. The dotted red line at July 2021 marks the presumed start of the SCTLD outbreak. The size of the dots corresponds to prevalence of tissue loss in individuals of a given species at a given site. The box represents the first and third quartiles and the horizontal line represents the median across sites. Letters indicate significant (p<0.05) differences among time points within each species.

### 3.3 SCTLD changed the composition of CBC reefs

We documented a significant change in benthic cover across major functional groups and over the survey period in photoquadrat data (GLMM survey timepoint x functional group, Chisq = 400.319, Df = 6, p <0.001; Fig 5). This change was driven by a substantial decrease in cover of living scleractinian corals across all study sites (Fig. 6, Table S4&S6) over time, from an average of 13.2% prior to the SCTLD outbreak, to 9.6% in May 2022, and 7.9% by December 2022, indicating a loss of 40.2% averaged across sites. CBC Lagoon, which had the highest scleractinian cover at all three timepoints, experienced a 42% decline from 26% to 15%. By December 2022, CBC30N had only 3% coral cover. Meanwhile, cover of turf algae increased (Fig. 5, Table S4) from 35.4% during the initial surveys, to 48.9% by December 2022. Octocoral cover declined slightly between 2019 and 2022, although there was no significant difference between May and December 2022 (Fig. 6, Table S6). This decrease was largely driven by a 30.4% decline at CBC Central and 33.8% decline at CBC30N. Other sites did not experience similar declines. Macroalgae cover largely stayed the same, with a significant but slight decrease in December 2022 (Fig. 6, Table S6).

**Figure 6.**
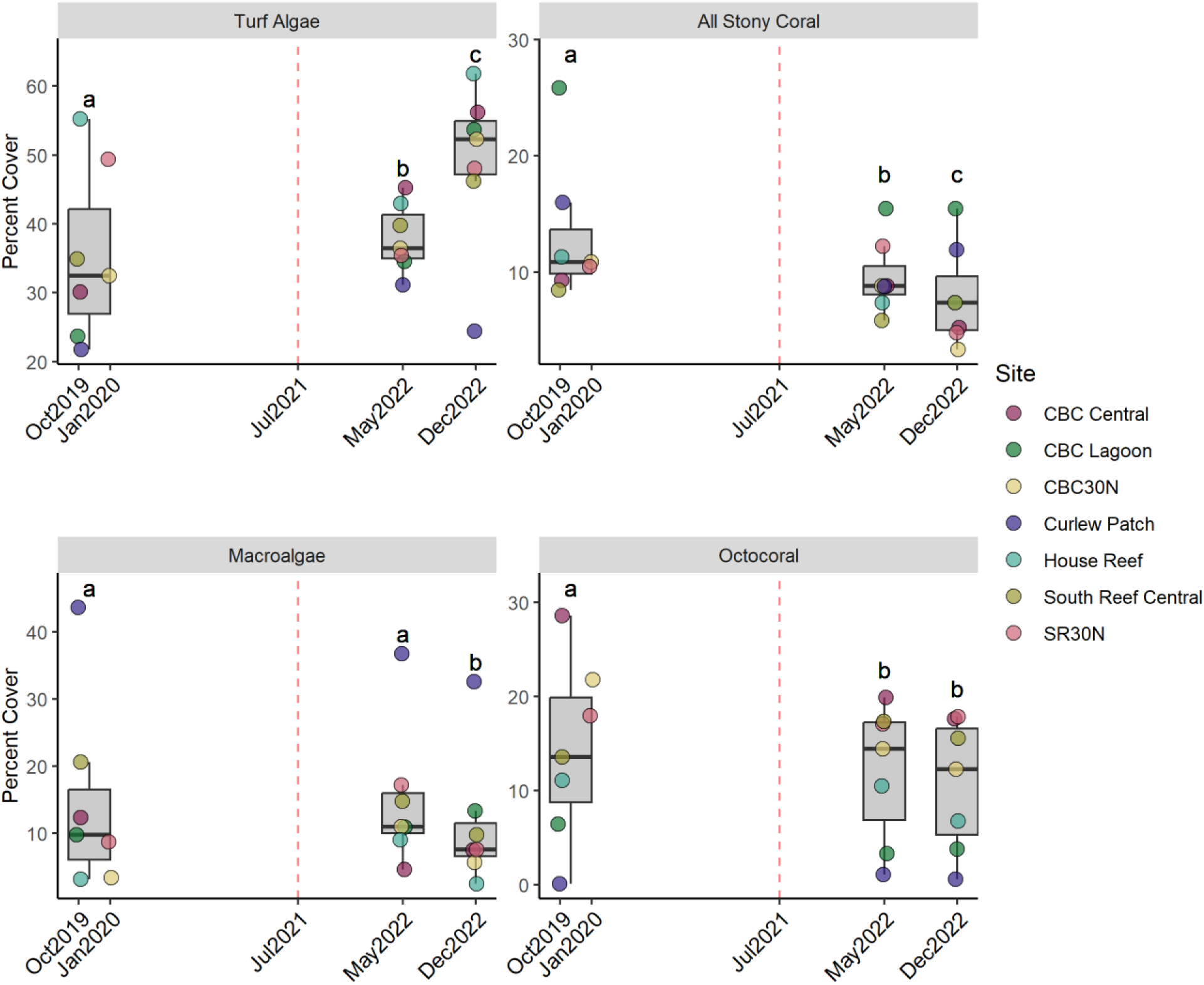
Percent cover of four benthic functional groups derived from photoquadrats taken along a 30×1 meter transect at each site over time. The dotted red line at July 2021 marks the presumed start of the SCTLD outbreak. The box represents the first and third quartiles and the horizontal line represents the median across sites. Letters indicate time points that differ significantly (p<0.05) within each functional group.

Within scleractinian corals, we found that the community composition changed over time. Using both photoquadrat and in situ survey data, we found that coral cover varied by species over time (GLMM photoquadrat species x survey timepoint, Chisq = 47.9, Df = 18, p < 0.001; GLMM in situ survey species x survey timepoint, Chisq = 46.993, Df = 26, p = 0.007). In addition, in situ community composition of adult corals >4cm changed significantly over time (PERMANOVA, Df = 2, SumOfSqs = 0.2056, R^2^ = 0.2501, F = 3.0012, p<0.001, Fig. S3). Due to limitations inherent to these two types of data as well as differences among coral species, we found the combination of these data produced a more complete picture of the impacts of SCTLD on coral community composition.

Confirming the results of tagged colony tracking, the in situ surveys revealed a total loss of *M. meandrites* (n = 8 in October 2019/January 2020) from within the survey transects by May 2022. Surveys also documented the total loss of *Eusmilia fastigiata* (n = 9 in October 2019/January 2020) by December 2022. At the end of the monitoring period, only one colony of *Diploria labyrinthiformis* and one colony of *Dichocoenia stokesii* remained (Fig. 5, significant decline over time, p=0.011). Because these species were relatively scarce at the start of this study and do not generally form large colonies, their declines were not captured in photoquadrat analyses of species-specific benthic cover change.

*Pseudodiploria strigosa* was found in survey data at all transects at the start of the study but declined in number at all transects and was absent from two by December 2022. Significant decreases were also observed between October 2019/January 2020 and May 2022 in *M. cavernosa*, and the *Orbicella* species complex (Table S3). Declines in the contribution of all three of these species to benthic cover were also observed in photoquadrat data between 2019 and May 2022, although each species covered less than 6% of the benthos at each site (Fig. 7, p < 0.03 for all, Table S5&S6).

**Figure 7.**
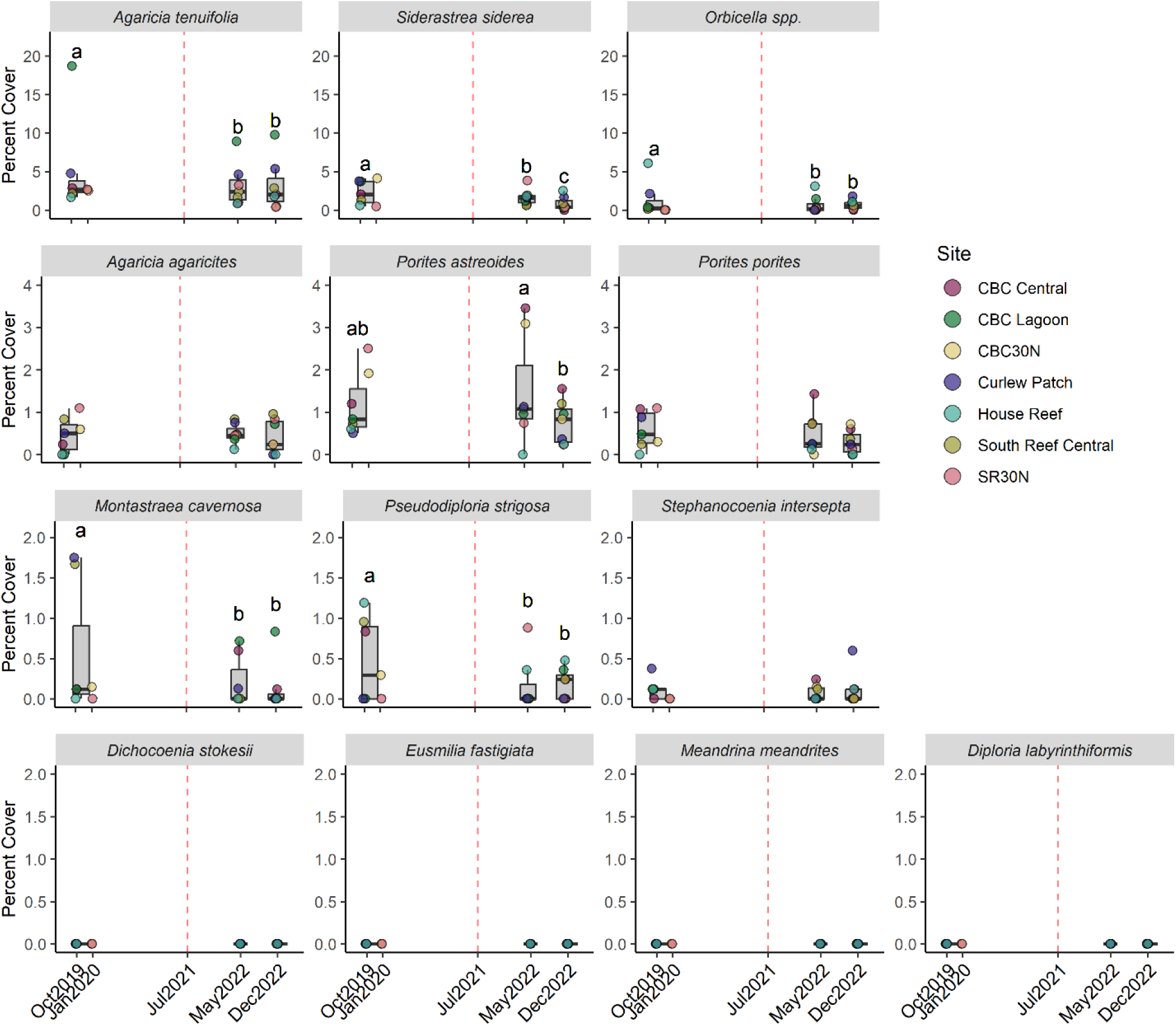
Percent cover of select species derived from photoquadrats taken along a 30 meter transect at each site over time. The dotted red line at July 2021 marks the presumed start of the SCTLD outbreak. The box represents the first and third quartiles and the horizontal line represents the median across sites. Letters indicate significant (p<0.05) differences among time points within each species.

*S. siderea* experienced a more gradual decline, from a total of 436 in 2019 (across all seven transects), to 159 in May 2022 (Fig. 5, Table S2&S3, p=0.006), but the number counted in December 2022 (235) did not differ significantly from the start. However, percent cover of *S. siderea* determined using photoquadrat data declined significantly between October 2019/January 2020 and May 2022 (Fig. 7, p < 0.03, Table S64) and again into December 2022 (p = 0.001).

*A. agaricites* and *P. porites* both declined in abundance according to in situ surveys from October 2019/January 2020 to May 2022 (Fig. 5, Table S3, p=0.001 and p=0.048, respectively) but their final abundances in Dec 2022 did not differ significantly from the start of the study period (Fig. 5) and a corresponding decrease in cover from photoquadrat data was not found. In contrast, cover of *A. tenuifolia* declined significantly only in photoquadrat data from the initial timepoints in October 2019/January 2020 to May of 2022 (Fig. 7, p < 0.001), but not December 2022. Cover of *P. astreoides* differed between May and December of 2022 (p = 0.002), but neither timepoint varied substantially from the initial photoquadrat data prior to the onset of SCTLD in the region and there was no corresponding decrease in colony count from in situ survey data.

## 4. Discussion

The initial outbreak of SCTLD had a profound effect on the health of coral reefs around Carrie Bow Cay, Belize. Our observed decline in both stony coral cover and number of colonies was mostly due to the loss of SCTLD-susceptible species, changing the species composition of these reefs. Tracking colonies through time allowed us to confirm the lethality of this disease as well as its variable impacts on different coral species not only in prevalence but also incidence and time to death.

While it requires significant time and effort, fate-tracking individual colonies can provide critical data about the onset of disease signs and the link between infection and mortality. Prior to this study, only four others have tracked the fates of initially healthy colonies through an SCTLD outbreak, all during the initial years of the disease in Miami-Dade county (Precht et al. 2016), the Middle Keys (Sharp et al. 2020), and the Upper Keys (Rippe et al. 2019; Kolodziej et al. 2021) of Florida. These studies used various fate-tracking methods - photomosaics (Kolodziej et al. 2021), established plots (Sharp et al. 2020), monitoring of tagged colonies (Precht et al. 2016), and skeletal coring of tagged colonies (Rippe et al. 2019) - to provide insights into species-specific disease prevalence and mortality during the initial SCTLD outbreak. The first published report of the disease (Precht et al. 2016) tracked 115 tagged colonies across 13 coral species and documented 100% mortality of all infected individuals across species.

While we likely missed peak prevalence of the disease across all susceptible species, we were still able to observe disease progression in several colonies of *M. cavernosa* and *P. strigosa* and document peak prevalence for *S. siderea*. As observed by others (Precht et al. 2018; Brandt et al. 2021; Aeby et al. 2025), the lesions we recorded in *S. siderea* largely differed in their distribution and rate of progression from those of other species, tending to be more diffuse and much slower to progress (Fig. 2, Fig. 4). Of course, it is impossible to be certain that the same pathogen(s) are causing the disease signs in *S. siderea* and other species as the causative agent of SCTLD remains unknown and diagnostic tools are limited (Papke et al. 2024). However, the fact that disease onset across most of our tagged *S. siderea* colonies occurred immediately after disease signs were observed in other species suggests they are likely at least related if not the same disease. Furthermore, Meiling et al. (2021) confirmed transmission of tissue loss signs from *D. labyrinthiformes* to *S. siderea* in the lab, and Aeby et al. (2025) confirmed transmission from *Colpophyllia natans* to *S. siderea*. Interestingly, *S. siderea* was not observed with disease signs in the early report by Precht et al. (2016) of the outbreak in Miami, although the later timing of disease onset in our tagged *S. siderea* colonies relative to other species suggests that the Miami *S. siderea* colonies may have developed lesions after the study concluded in 2015. Rippe et al. (2019) also observed a slower onset of disease and disease progression in *S. siderea*affected by SCTLD.

Across species, we observed high mortality during the study period, further solidifying SCTLD as one of the most deadly coral diseases ever reported (out of the 102 colonies tagged in 2019, 42% were dead by December). As reported in other locations, mortality and timing varied greatly across species and largely followed previously reported patterns. The 100% total mortality before May 2022 that we report in *M. meandrites* is comparable to the 100% mortality of this species observed by Precht et al. (2016) and the near complete mortality reported by Sharp et al. (2020), although the one *M. meandrites* observed by Kolodziej et al (2021) was still alive at the end of the study. Our results add to the growing body of literature documenting this species as one of the first to show disease signs and quickest to succumb to the disease (Papke et al. 2024). Monitoring efforts for SCTLD in areas still unaffected by the disease will likely be most fruitful if concentrated on this species especially given its relatively large size when compared to other highly susceptible species (e.g., *Eusmilia fastigiata* and *Dichoecoenia stokesii*). *P. strigosa* experienced the next most rapid partial and whole colony mortality, with nearly 50% of colonies dying prior to the May 2022 observation at which point two had disease signs, which were then dead by December. The 71% whole-colony mortality we observed in this species by Dec 2022 was similar to the 68% total mortality reported by Kolodziej et al. (2021) across a slightly shorter timeframe and the 79% mortality reported in Sharp et al. 2020. Rippe et al. 2019 similarly observed rapid colony mortality in *P. strigosa* within a year after onset of disease in the area. They did not, however, observe any disease signs in *P. strigosa* at inshore reef sites during this time period, leading to a lower percent mortality (30%) across both sites. Our results suggest a much slower rate of mortality due to SCTLD for most *M. cavernosa*: only 29% were dead before May 2022 and two of the seven colonies with disease signs in May 2022 were still alive and diseased in December, and one had no active tissue loss. Sharp et al. (2020) also observed lower total mortality (25%) in *M. cavernosa,* and Kolodziej et al. (2021) observed the lowest with 8% total mortality. This slow overall progression may be reflective of an initial high mortality of susceptible *M. cavernosa* colonies during the onset of disease followed by slower mortality as the disease becomes endemic (Aeby et al. 2019, 2021). Across all species, only 21% of the colonies we observed with lesions in May 2022 were dead by that December, in contrast to the 100% mortality reported by Precht et al. (2016) one year after peak prevalence during the initial outbreak. This discrepancy is likely largely due to the fact that Precht et al. did not document any *S. siderea* with disease signs during their study period. In fact, in our study, the majority of the corals that were lesioned but did not die during the survey period belonged to *S. siderea* (18 out of the 21 diseased in May 2022 were still alive and diseased in December). Other studies similarly saw slower onset of disease and less or delayed whole-colony mortality in *S. siderea* colonies (Rippe et al. 2019; Sharp et al. 2020; Kolodziej et al. 2021). However, even limiting our analysis to *M. cavernosa* and *P. strigosa*, we did not see 100% mortality by the end of our study period, potentially suggesting a change in virulence or disease progression between this version of the disease and the initial outbreak (Aeby et al. 2021). We observed no mortality of *P. astreoides*, which is generally considered to have low susceptibility to SCTLD (Aeby et al. 2019; Sharp et al. 2020; Brandt et al. 2021; Alvarez-Filip et al. 2022), and in fact two of the three colonies with signs of tissue loss in May 2022 had no active tissue loss in December, suggesting these colonies may have been experiencing tissue loss due to other causes or recovered from SCTLD before complete mortality occurred. Sharp et al. (2020) also observed a few *P. astreoides* colonies with inactive tissue loss following SCTLD disease signs. Experimental transmission of SCTLD to *P. astreoides* has been observed in one study in the USVI (Meiling et al. 2021), but not another in Florida (Aeby et al. 2019), so the susceptibility of *P. astreoides* is still unclear or may vary regionally.

Monitoring for in situ lesion prevalence and recent mortality along fixed or random transects or in roving diver surveys has been a more common method than fate-tracking of tagged colonies to estimate SCTLD prevalence, especially in the Mesoamerican Barrier Reef regions that were exposed to the disease prior to Carrie Bow Cay (Alvarez-Filip et al. 2019, 2022; Estrada-Saldívar et al. 2020; Lee Hing et al. 2022). However, directly comparing lesion prevalences among studies that surveyed sites at single time points is challenging because relative time between the survey and peak prevalence is not standardized and in many cases unknown. Still, evaluating the relative susceptibilities of species is useful for identifying geographic variation in disease patterns. Even with multiple timepoints, we missed peak lesion prevalence among the most susceptible species. While our transects initially had low numbers of these species, all *D. cylindrus* and *M. meandrites* and most *E. fastigiata* and *D. stokesii* were already absent from transects by our initial post-outbreak surveys in May 2022 (Table 1, Fig. 5). This rate of mortality is consistent with the relative susceptibilities reported elsewhere in the region (Alvarez-Filip et al. 2019, 2022; Estrada-Saldívar et al. 2020; Heres et al. 2021) and with the results from our tagged *M. meandrites*. Similarly, survey data corroborated the results of our tagged colonies, with the highest prevalence in *P. strigosa* in May 2022 and lower prevalences in *M. cavernosa* and *Orbicella* spp. at both timepoints. Survey data of *S. siderea* found prevalence increased between May and December 2022, further supporting our findings from tagged colonies that disease progression is slower in this species.

Notably, in the only other peer-reviewed report from Belize thus far (although at reefs further north than CBC), Lee Hing et al. (2022) reported substantially lower disease prevalences for many susceptible species than what we observed in our transect surveys. This discrepancy is likely due to survey timing: Lee Hing et al. performed rapid response assessments very early in the disease outbreak (within a week of first report in Honduras in October 2020), while in this study we report lesion prevalence and mortality nearly a year after the estimated arrival of SCTLD at Carrie Bow Cay. Further supporting this claim, they were able to record 68% prevalence of disease signs in *M. meandrites* suggesting that prevalence likely increased in species like *P. strigosa* and *M. cavernosa* in subsequent months.

Cover of all scleractinian corals, the foremost reported indicator of coral ecosystem health (Acosta-Chaparro et al. 2025), declined significantly during our study primarily due to SCTLD. Relative loss of coral cover due to SCTLD outbreaks has varied across Caribbean locations from as low as 1.7% (Cheeca Rocks, Upper FL Keys, (Kolodziej et al. 2021)) to as high as 67% (inner reef terrace of coastal south FL, (Jones et al. 2022)), with relative losses of approximately 40-60% reported in Puerto Rico (Williams et al. 2021), Cozumel, Mexico (Estrada-Saldívar et al. 2020), St Thomas, USVI (Brandt et al. 2021), Turks and Caicos (Heres et al. 2021), and Bonaire (Pepe et al. 2025). Average coral cover around Carrie Bow Cay declined by approximately 40% during the course of monitoring, consistent with loss severities reported throughout the Caribbean. Variability in cover loss between locations likely is due to a combination of factors including starting cover and density of colonies, species composition, variation in monitoring duration, and disease severity. Even before the onset of SCTLD, Belize’s barrier reef had been losing coral cover and shifting in composition (Alves et al. 2022), though recent spatially-limited periods of recovery have been reported near Carrie Bow Cay (De Pablo et al. 2021; Mumby et al. 2021). The transects monitored here were originally dominated by *A. tenuifolia*, *S. siderea*, and *Orbicella* spp in terms of species-specific scleractinian cover (not density or absolute number of colonies), but with the decline of those species, weedy *P. astreoides* colonies emerged as a co-equal cover contributor by the end of the study period. Our prevalence and fate tracking data confirm that loss of *Orbicella* spp. and *S. siderea* were due to SCTLD. However, *A. tenuifolia* also experienced cover declines despite its relatively low SCTLD susceptibility, especially at one site (CBC Lagoon), (Fig. 7). We suspect the declines in this weedy, physically fragile species at this shallow patch reef were the result of Hurricane Nana, which affected the area in September 2020. While unlikely, since direct effects of the hurricane (or disease) were not documented due to the closure of the research station, disease could have potentially played a role in the decline as the susceptibility of *A. tenuifolia* to SCTLD remains poorly described (Papke et al. 2024), and other Agaricia species have been documented with acute tissue loss in epidemic regions (Alvarez-Filip et al. 2019; Costa et al. 2021). Many species that are highly susceptible to SCTLD do not contribute significantly to coral cover as measured through top-down benthic photoquadrats due to their small sizes and/or cryptic microhabitat preferences (Costa et al. 2021), underscoring the importance of monitoring for change in reef ecosystems using multiple metrics.

Colony density is a critical variable to measure throughout mass coral mortality events because as slow-growing sessile organisms, corals are particularly vulnerable to Allee effects. In an Indo-Pacific broadcast-spawning acroporid, recent research has revealed that the 30% fertilization success between closely situated (<0.5m) colonies drops to 10% when nearest neighbors are separated by just 10m (Mumby et al. 2024). Here, we report significant declines in the number of colonies counted in a 30m^2^ area for three species (*M. cavernosa*, *Orbicella* spp., and *D. stokesii*), as well as total loss of *M. meandrites* and *E. fastigiata* from survey transects (Fig. 5). While some highly abundant species (*A. agaricites*, *P. porites*, *S. siderea*) varied significantly between the initial surveys and May 2022 but rebounded in abundance by December 2022, this pattern may be partially attributable to high temporal variation in recruitment and thus in abundances of small colonies (Harper et al. 2023), and in the case of the branching *P. porites* complex, the result of temporal variation in colony breakage and separation. Determining conclusively whether the declines in abundance on our transects are representative of density declines throughout the surrounding reef system, and whether those losses will precipitate Allee effects that inhibit recruitment of SCTLD-susceptible species in the future, will require synthesizing monitoring data at the regional scale as well as continued tracking of recruitment in situ into the future.

Significant shifts in community assemblage have been reported throughout the Caribbean and Florida in the wake of SCTLD (Alvarez-Filip et al. 2019; Brandt et al. 2021; Croquer et al. 2022; Hayes et al. 2022b; Jones et al. 2022). Likewise, we observed a marked shift in scleractinian community composition between October 2019 and December 2022 at the reefs around Carrie Bow Cay (Fig S3). As has been observed in other locations, our transects became more dominated by weedy or stress-tolerant agariciids, poritids, and siderastreids (Darling *et al*., 2012). This shift underscores the hypothesis that SCTLD is further reducing the functionality of Caribbean coral reef ecosystems by reducing populations of many large boulder coral species and flattening both structural and taxonomic complexity through a reduction in coral cover, an increase in turf algae cover, and a weedier coral assemblage comprised of mostly smaller-growing, shorter-lived species (Alvarez-Filip et al. 2019, 2022). SCTLD may affect reefs with higher coral species diversity (Williams et al. 2020; Muller et al. 2020), contradicting the diversity-disease hypothesis in which high host diversity buffers communities against disease risk (Keesing et al. 2006; Muller et al. 2020; Costa et al. 2021; Pagenkopp Lohan et al. 2024). This contradiction likely occurs because rare species exhibit high susceptibilities, as shown here with *M. meandrina* and *E. fastigiata*, but many abundant species are intermediately susceptible and subject to disease transmission, giving SCTLD the potential to devastate diversity, cover, and functionality across Caribbean reefs.

With increasing global sea temperatures, prevalence of coral diseases has increased across the globe (Randall 2014; Tracy et al. 2019; Burke et al. 2023)(, particularly in the Caribbean which has historically been a hotspot of coral disease (Morais et al. 2022). Our study supports previous literature that SCTLD is one of the deadliest coral diseases seen to date due to high infection and mortality rates (up to 100%) affecting a large proportion of species. Its rapid progression through the Caribbean warrants an immediate need for research and interventions. Continued monitoring over larger spatial and temporal scales than those reported here is necessary to fully understand the scope of the ecosystem degradation caused by SCTLD in Belize and to plan for mitigation strategies that support future disease and disturbance resilience. While various interventions, including culling, treatment with antibiotics and probiotics, and genetic rescue have been implemented throughout the Caribbean (Neely et al. 2021b; Papke et al. 2024), the Belize Barrier Reef is vast and largely remote, and existing interventions are challenging, expensive, or impossible to scale.

## Acknowledgements

This research was funded by an NSF RAPID # 2347450 and an award from the Paul M. Angell Family Foundation to SGW, an NSF GRFP to BKS, a Smithsonian Life on a Sustainable Planet award to LH and SGW with additional support from the Smithsonian MarineGEO program. We would like to acknowledge Branae Craveiro and Caroline DeSouza for help organizing photos and associated data as well as Lucia Rodriguez, Sara Swaminathan, Ximena Boza, and Kevin and Earl David for help with fieldwork. This is contribution #XXX from the MarineGEO network.

