## Supplemental Figures for "Stony coral tissue loss disease outbreak altered reef communities of Carrie Bow Cay, Belize"

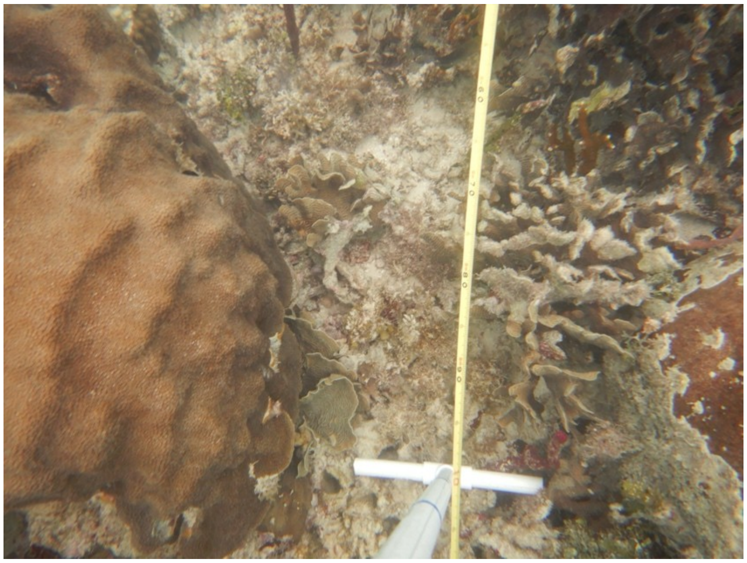

Figure S1. Example photoquadrat.

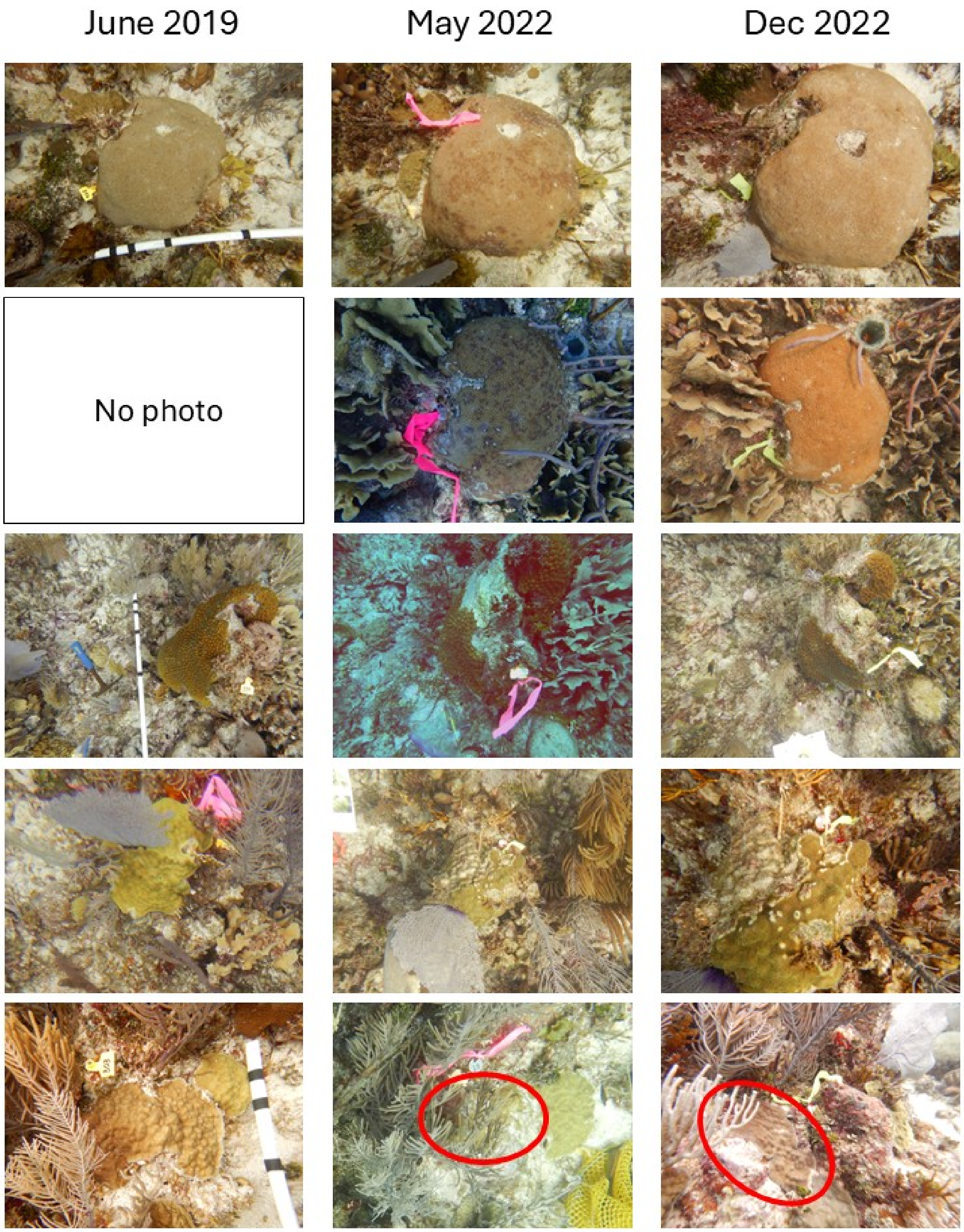


Fig. S2 Recovery in five individuals (colonies that were sampled as diseased in May 2022 but appeared healthy by December 2022).

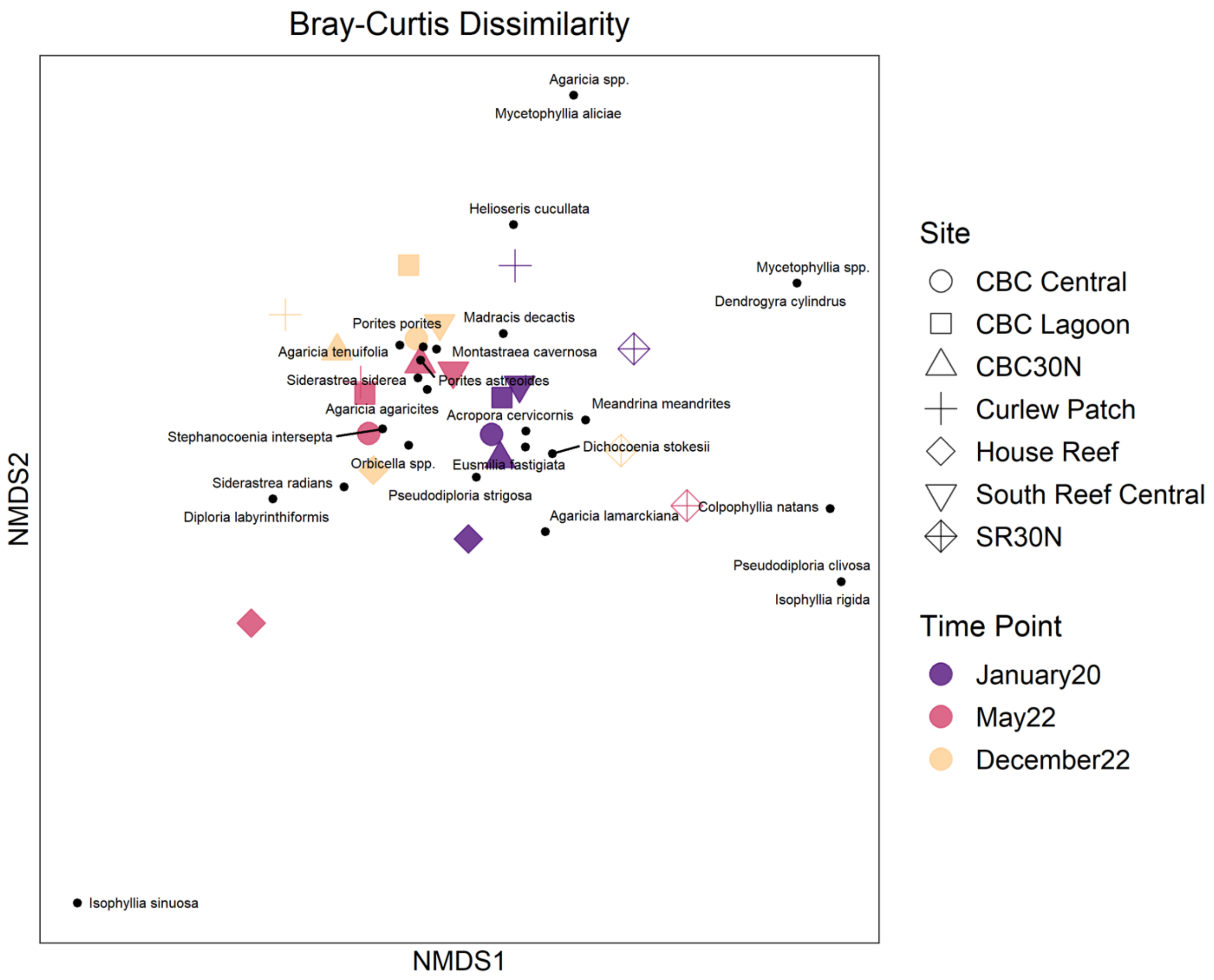


Fig. S3. nMDS of in situ coral community composition for colonies >4cm.
